# Universal Features of Epidemic Models Under Social Distancing Guidelines

**DOI:** 10.1101/2020.06.21.163931

**Authors:** Mahdiar Sadeghi, James M. Greene, Eduardo D. Sontag

## Abstract

Social distancing as a form of nonpharmaceutical intervention has been enacted in many countries as a form of mitigating the spread of COVID-19. There has been a large interest in mathematical modeling to aid in the prediction of both the total infected population and virus-related deaths, as well as to aid government agencies in decision making. As the virus continues to spread, there are both economic and sociological incentives to minimize time spent with strict distancing mandates enforced, and/or to adopt periodically relaxed distancing protocols, which allow for scheduled economic activity. The main objective of this study is to reduce the disease burden in a population, here measured as the peak of the infected population, while simultaneously minimizing the length of time the population is socially distanced, utilizing both a single period of social distancing as well as periodic relaxation. We derive a linear relationship among the optimal start time and duration of a single interval of social distancing from an approximation of the classic epidemic *SIR* model. Furthermore, we see a sharp phase transition region in start times for a single pulse of distancing, where the peak of the infected population changes rapidly; notably, this transition occurs well *before* one would intuitively expect. By numerical investigation of more sophisticated epidemiological models designed specifically to describe the COVID-19 pandemic, we see that all share remarkably similar dynamic characteristics when contact rates are subject to periodic or one-shot changes, and hence lead us to conclude that these features are *universal* in epidemic models. On the other hand, the nonlinearity of epidemic models leads to non-monotone behavior of the peak of infected population under periodic relaxation of social distancing policies. This observation led us to hypothesize that an additional single interval social distancing at a *proper time* can significantly decrease the infected peak of periodic policies, and we verified this improvement numerically. While synchronous quarantine and social distancing mandates across populations effectively minimize the spread of an epidemic over the world, relaxation decisions should not be enacted at the same time for different populations.

## 1 Introduction

COVID-19, the disease first identified in Wuhan, China and the cause of the 2020 pandemic, has infected over 90 million people worldwide, caused at least 1.9 million deaths [1], and has resulted in a worldwide economic downturn [2]. After the shelter-in-place ordinances [3], social distancing as a form of Non-Pharmaceutical Intervention (NPI) has been enacted in the United States [4], and other countries [5, 6] for reducing the spread of the virus, as neither herd immunity nor a viable vaccine yet existed [7]. Many countries have implemented strict quarantine, isolation, or social distancing policies early in the epidemic [5], while countries such as Belarus [8] and Sweden [9,10] have taken more lenient approaches at the onset of the outbreak. Understanding optimal strategies for social distancing will both “flatten the curve” and hopefully ease the economic burden experienced due to prolonged economic stagnation [11–13]. The goal of this manuscript is thus to investigate the response of the disease to different time-varying social distancing strategies.

### 1.1 COVID-19 and social distancing

There has been much recent theoretical work revisiting, expanding, and studying dynamical and control properties of classical epidemic models so as to understand the spread of COVID-19 during quarantine and social distancing [14–19], including studies of (integral) input to state stability [20], network stability of epidemic spread [21, 22], and optimal control strategies for meta-population models [23]. These models have been used to predict the potential number of infected individuals and virus-related deaths, as well as to aid government agencies in decision making [24]. Most models are variations on the classical Susceptible-Infected-Recovered (*SIR*) model [25–27] which have been modified to more closely predict the spread of COVID-19. Some such extensions are listed below:

1. Expanding the *SIR* model to include additional population compartments. Such compartments may describe individuals that are placed under quarantine and/or in social isolation. Other models explicitly subdivide populations into both symptomatic and asymptomatic infected individuals [28–33], as it is currently thought that COVID-19 is significantly spread through *asymptomatic* individuals [34–36].
2. Modeling the effects of social distancing for an infection aware population. This can be done by changing the contact rates between the compartments, or by modeling the behavior of a population that alters its social interactions because of observed infections or deaths [37, 38]. The latter technique has recently been applied to COVID-19 [39, 40].
3. Sub-dividing populations into regions, each described by *local* parameters. Such regions may be cities, neighborhoods, or communities [41]. This framework allows modelers to capture the virus spread and population mobility geographically [42–45]. These models have been recently used to understand the spread of COVID-19 in China [46], Italy [47], Netherlands and Belgium [48], and India [49, 50].

Shortening the period of time that populations are socially distanced is economically advantageous [2, 51, 52]. The main objective of this study is to reduce the disease burden (here measured as the peak of the infected population) while simultaneously minimizing the length of time that the population is socially distanced.

### 1.2 Periodic relaxation policies

The timing of one-shot pulses is analyzed in this work, but other policies may be preferred economically. For example, we also consider periodically relaxing social distancing mandates, which allow for scheduled economic activity. We find that the peak of the infected compartment is non-monotonic with respect to both the scheduling period and the ratio between the time of social distancing and the period [28, 53]. Such strategies can delay the infected peak. Our simulations and analysis suggest that the peaks of infected populations under periodic relaxation policies can be significantly reduced (between 30 – 50%), if they are combined with a strict single pulse of social distancing at a *proper time*.

### 1.3 Outline of this work

In the following section, we begin by giving a mathematical description for one-shot social distancing policy during an epidemic. We investigate the overall peak reduction in the infected population of the classic *SIR* model and a number of its variants in Section 3. We numerically discover a near linear optimal trade-off between the social distancing start time and duration, which seems to be ubiquitous among all models considered. This behavior can be explained by analytical considerations, at least for the *SIR* model. In Section 4, we summarize the conclusions as well as limitations of the work presented.

## 2 Mathematical modeling framework

Deterministic population models are commonly used to model the spread of an epidemic like COVID-19 [25, 26, 28]. Although some assumptions (such as population homogeneity) are not precisely accurate, such frameworks still provide invaluable insight in predicting the spread of the disease, and can be utilized to inform policy decisions in the presence of a pandemic.

We note that in epidemic models described by Ordinary Differential Equations (ODEs) where the recovered population is immune to reinfection (no transitions from the recovered compartment back to the susceptible compartment), infections may be eradicated *only asymptotically* (see Appendix D); that is, at any finite time, the proportion of infected individuals will be non-zero. Of course, physically this is unrealistic, and if the proportion of individuals affected is less than the proportion of a single person, the disease should be deemed eradicated. We note that this can be implemented via a threshold for the infected compartment (e.g. a proportion of 10^−7^), where the disease is considered eradicated once the infected compartment drops below this threshold. Alternatively, one may use corresponding stochastic models that may (with non-zero probability) reach zero in finite time (e.g. [33]).

Obviously, a strict social distancing or quarantine mandate at the very beginning of an epidemic may significantly impede the infection. In this case the number of remaining susceptible population (*S_t→∞_*) will be close to the initial susceptible population (*S*_*t*=0_). In other words, the number of total infected individuals will be limited to *I*_*t*=0_. However, this might be difficult to achieve for highly contagious diseases such as COVID-19.

### 2.1 Review of *SIR* model

As a first step, before considering more complex scenarios, we review the standard *SIR* model [25]. This model takes the form of a set of three coupled ODEs as follows:

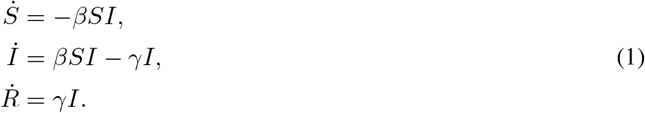

Here *β* is the transmission rate between the susceptible *S* and infected *I* individuals in a well-mixed population, and *γ* is the rate of recovery into the recovered (removed, here immune) *R* compartment. The *SIR* model can be used as an approximation to COVID-19 dynamics if immunity is long-lasting, which seems appropriate given time scales of the distancing policies considered here (less than one year, see below). Note that deceased and recovered individuals are combined into a single removed compartment *R*. The initial conditions are fixed as (*S*(0), *I*(0), 0) with a small initial value for the infected compartment *I*(0), and

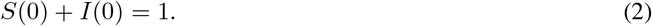

That is, we have normalized the variables so that all compartments denote percentages of the population, and not raw population numbers. We may translate directly to raw population numbers by multiplying the fractions (*S, I, R*) by the initial susceptible population *S*(0).

### 2.2 Single interval social distancing

Social distancing is currently being implemented via a variety of different techniques. For example, in many locations local and federal governments are issuing rules related to quarantine and isolation, as well as regulations for wearing masks and avoiding non-essential interactions to reduce contact rates [54]. In the *SIR* model discussed in Section 2.1, social distancing may be mathematically modeled by temporarily reducing the transmission rate of the disease, i.e. parameter *β*.

As an example, consider a single interval of social distancing (a one-shot pulse), represented mathematically as a time-varying transmission rate *β*(*t*) in Eq. (3):

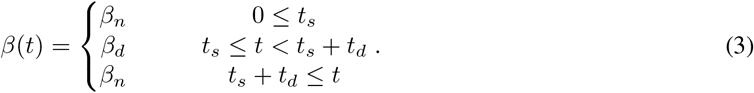

Here we assume that *β* can be effectively reduced from *β_n_* (contact rate during normal time for non-distanced population) to *β_d_* (contact rate during social distancing) during distancing. Here distancing regulations are enacted at day *t_s_*, which corresponds to a decrease in the transmission rate *β_n_* → *β_d_*. Regulations are implemented until day *t_s_ + t_d_*, after which they are completely relaxed, i.e. *β_d_* → *β_n_*. Note that for other models, distancing may be implemented differently. For example, in an infection-aware community, viral transmission may be reduced as the number of confirmed cases increases during an epidemic [39], hence yielding a feedback law governing *β = β*(*I*). For a visualization of (3), see Fig. 1.

**Figure 1:**
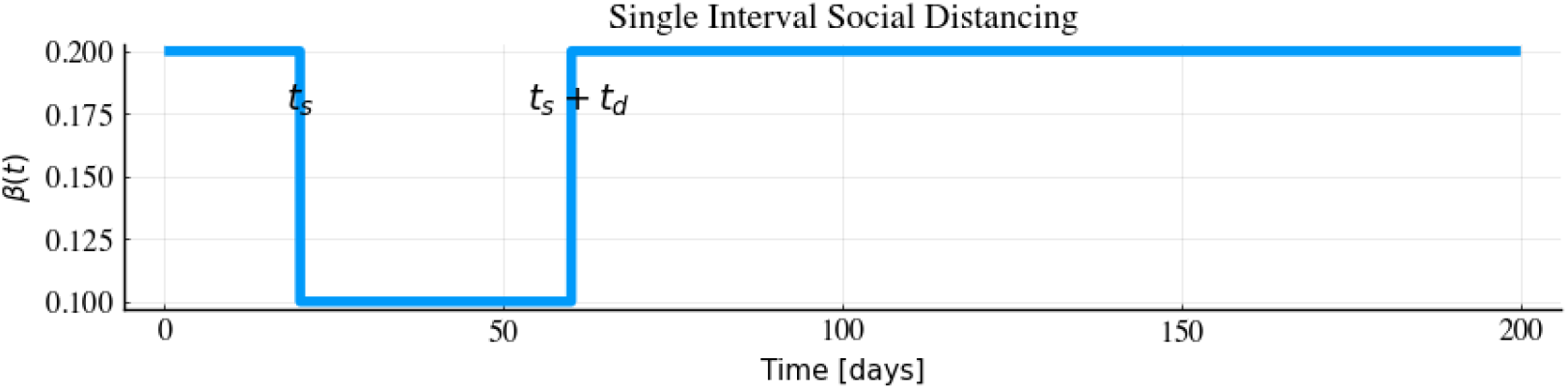
Single interval social distancing during an epidemic for the *SIR* model. Here social distancing starts at day *t_s_* = 20, and lasts for *t_d_* = 40 days, after which regulations are completely relaxed.

We investigate how the start time *t_s_* and duration *t_d_* affect the peak of the infected compartment. Note that prolonged distancing cannot be enacted without severe economical/sociological consequences, so that the timing of an *interval* of distancing is of great interest. A rigorous analysis has been performed in [55] to find the optimal schedule of distancing based on the *SIR* model and cost function that minimizes a combination of the total number of deaths and the peak of the infected compartment. Indeed, the authors in [55] find a distancing policy of the form (3) as optimal, which motivates us to investigate the precise switching times *t_s_* and *t_d_*. That is, we are interested in understanding *when* social distancing should be implemented, hence called *properly timed* social distancing here, for a specific distancing period. In the following we will study this problem for a subset of recently proposed models that have been introduced to understand the spread of COVID-19.

## 3 Results

In this section, we discover an almost linear relationship between the start time *t_s_* and a finite duration of optimal one-shot social distancing *t_d_* (see Fig. 1, which represents a simple model of single interval social distancing as a one-shot pulse input). We begin with the *SIR* model and observe a linear trade off between the start time and duration of social distancing. We note a distinctive “V” shaped pattern in the heat maps/contour plots, and analyze this geometry mathematically. Then, we perform similar numerical simulations for other epidemic models that have been recently proposed for describing COVID-19. We find that the “V” shaped pattern is consistent among all of these models, and we believe it is a universal feature in epidemic models.

Lastly, we investigate periodic relaxation policies that have been recently proposed by [53, 56]; our goal is to understand their limitations. We also show that a well-timed one-shot social distancing pulse can significantly reduce the infected peak, *when applied in conjunction with periodic relaxation strategies*.

### 3.1 Single-pulse response

Fig. 2 represents the peak of the infected population in response to a single interval of social distancing for the *SIR* model (1). Fig. 2a represents the maximum of the infected population,

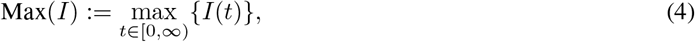

as a function of the distancing start time *t_s_* and duration *t_d_*. Note that dark blue color corresponds to high maximum infection, with lighter colors representing lower values in the peak of the infection. Based on the model parameters and initial conditions defined in Appendix C.1, it can be observed that the maximum infection has been reduced by 50% with properly timed distancing. For example: social distancing at disease onset (*t_s_* = 0) for a duration of *t_d_* = 50 days would not be effective in reducing maximum infection (Max(*I*) = 0.4) as the infected peak happens after the social distancing interval; however, by increasing the length of social distancing to 160 days (*t_d_* = 160), we observe a reduction in the maximum infection by more than 50% (Max(*I*) = 0.15). Note that the latter regimen also avoids a second wave of infection after the social distancing interval. Fig. 2 represents the infected peak Max(*I*), and *not* the time *t_p_* at which this peak occurs:

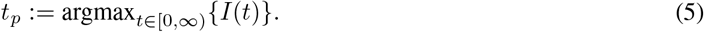

**Figure 2:**
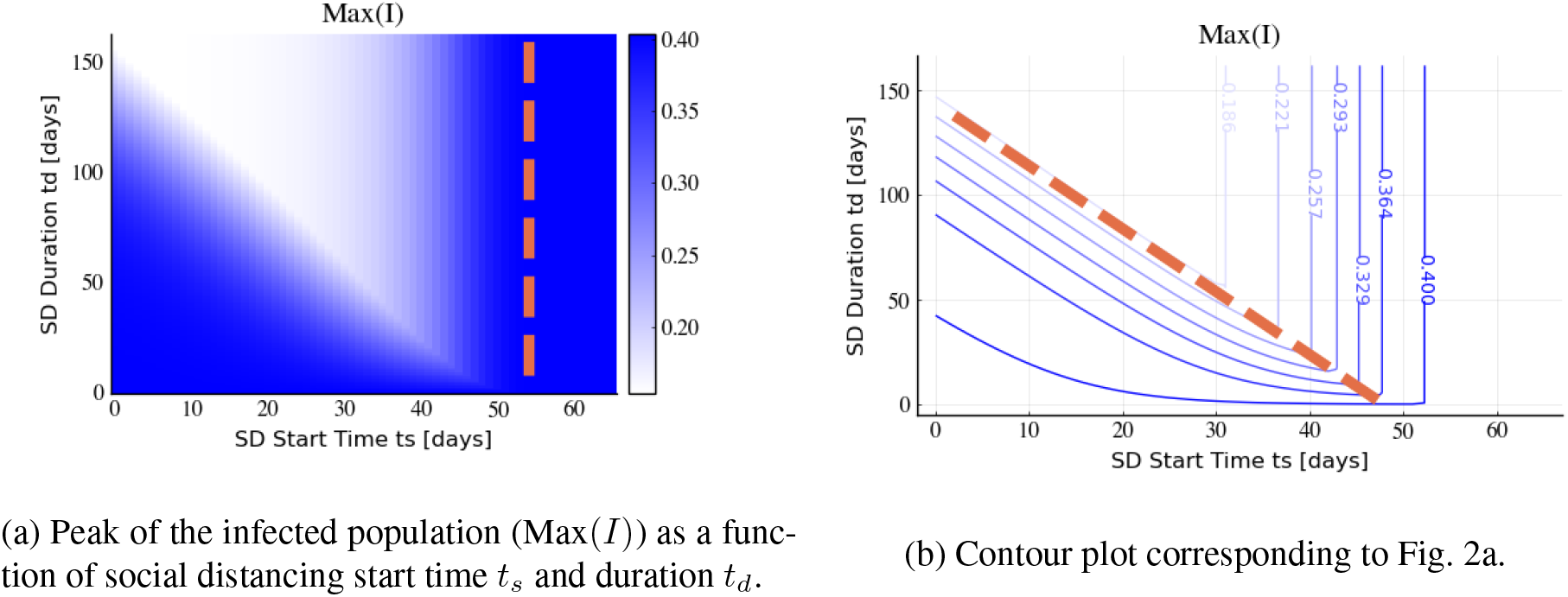
Infected peak, Max(*I*), under single interval social distancing (SD) policy. Time is measured in days, and the parameters used for the *SIR* model are *β_n_* = 0.2, *β_d_* = 0.1, and *γ* = 0.05 (see Eqs. (1) and (3)).

Similar representations as in Fig. 2 are used for other models considered in latter sections of the manuscript. For information on parameters used in this and all subsequent sections, see Appendix C.

The dashed vertical line in Fig. 2a indicates the time at which the infected peak under no distancing mandates would peak; we call this the “virtual peak.” Note that implementing social distancing after the vertical line (a *too late* start time *t_s_*) has no effect on infected peak reduction. In general, we observe that there are schedules such that one-shot pulses effectively reduce the infected peak up to 50% of the virtual peak. Specifically, from the left side of the dashed vertical line, we see that social distancing can be effective if it is implemented early enough.

We denote the combination of diagonal and vertical sections of the contours in Fig. 2b as a characteristic “V” shaped pattern. Intuitively, we expect a vertical line as the boundary between the blue and white area in Fig. 2a, because implementing *too late* means that distancing policies will have little effect on reducing the spread of the disease; there are already too many infected individuals. Indeed, it is clear that as we decrease *β* (enact social distancing), the peak of infected population decreases, *if social distancing is enacted quickly enough*. Similarly, any social distancing policy initiated after the virtual peak would be ineffective in decreasing the peak (see Section 3.2).

However, we note a large transition region in start times *t_s_ before* the non-distanced peak actually occurs; that is, distancing must be implemented before the occurrence of the peak of the infected population under no social distancing mandate, if we are to significantly inhibit the disease burden. This is non-intuitive, and it is important for policy-makers to understand this *gap* during which one-shot pulses are sensitive to the start time *t_s_*. This can be explained by sensitivity of the infected peak to start time, with details provided in Section 3.2. The infected peak is very sensitive to start times in a narrow band, due primarily to sensitivity of the integral curves of the SIR model in a region of the (*S, I*) phase-space. In the provided simulations (Fig. 2), the time scales are on the orders of days, but in general depend on specific parameter values.

The diagonal dashed line in Fig. 2b represents an almost linear trade-off between the start and the duration of the social distancing. We can understand this theoretically: we show below that the slope of this line is given approximately, as a function of the non-distanced and distanced transition rates and the removal (recovery) rate (*β_n_ β_d_*, and *γ* respectively), by the following formula:

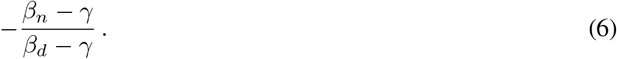

For example, the slope of the diagonal line in Fig. 2b is approximately − 3, with *β_d_* = 0.1, *β_n_* = 0.2, and *γ* = 0.05.

To derive (6), consider the linear approximation for the infected population (42), derived in Appendix A.2. For convenience, we provide the formula below:

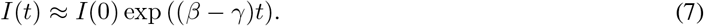

Hence, if *t_p_* denotes the time of the peak infected population, we have

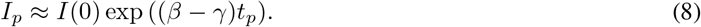

To understand the dynamics, intuitively one can utilize the above formula from *t* = 0 to *t* = *t_s_* with no social distancing enacted, and a one-shot pulse with duration of *t_d_* (i.e. from time *t* = *t_s_* to *t* = *t_s_* + *t_d_*). Note that the two formulas are connected via their initial conditions, which are *I*(0) and *I*(*t_s_*), respectively. Hence, the infected compartment at time *t = t_s_ + t_d_* can be approximated as

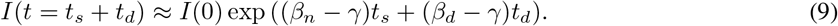

If the infected compartment reaches its fixed maximum value at time *t_p_ = t_s_ + t_d_*, then Eq. (9) implies that *t_s_* and *t_d_* are related by

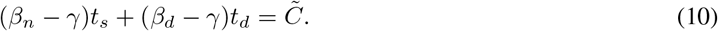

for a constant 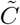. Here 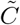 is a function of *I_p_*, but not *t_s_* and *t_d_*.

Hence the contours of constant *I*_max_ in Fig. 2b can be approximated by parallel lines, each with slope given by (6). By observing the contours of Fig. 2b, we see a similar parallel structure. The diagonal dashed line represents the slope (6), which appears to agree with the contour lines.

The blue region at the bottom left side of Fig. 2a represents the social distancing policies with a short duration that start *too early*. The higher infected peak indicates that the peak of the infected compartment occurs after distancing relaxation; note that such policies have approximately the same infected peak as if distancing was never implemented (compare to *t_d_* = 0 days). Although such policies delay the infected peak and provide more time for the discovery of new testing and treatment methods, they are not effective in reducing the infected peak.

Although it is not the main focus of this manuscript, we note that social distancing policies can be modeled in combination with mask mandates through the transmission rate *β* in the *SIR* model (and hence by extension, more complicated epidemic models). In general, *β* can be interpreted as a product of two terms: (a) the rate of meeting infected individuals, and (b) the probability of contracting the disease once an infected individual is encountered. Wearing masks reduces the latter, but does not inherently affect the former. Hence, when a specific social distancing protocol is implemented, the ratio between *β_n_* and *β_d_* is preserved between distancing and relaxation scenarios, as wearing masks reduces the rates *β_n_* and *β_d_* by the same constant factor. This provides a simple yet intuitive means of modeling the combined effects of mask mandates and social distancing, and it also allows us to predict how the “V” shaped pattern seen in Fig. 2 changes if masks are worn. An example is shown in Fig. 3, where the ratio between *β_n_* and *β_d_* remains as in Fig. 2, but smaller individual transmission rates are assumed due to individuals wearing masks. Note that from (6), a slightly different characteristic slope appears, but if *γ* is small, then this slope is approximately the same as in the non-masked case.

**Figure 3:**
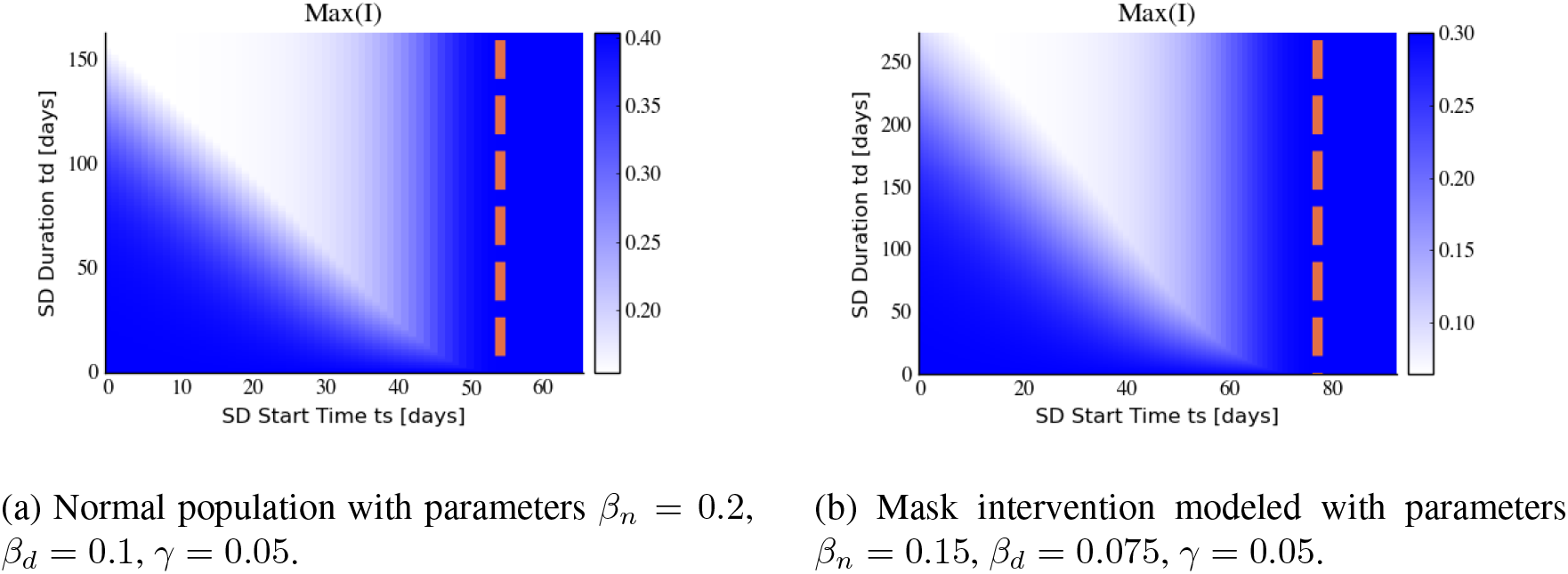
Infected peak, Max(*I*), for a single interval of social distancing for both a normal ((a)) and masked ((b)) population in *SIR* model (1). Both populations have the same ratio of transmission rates *β_n_*/*β_d_* = 2. The ranges of the plots are picked so as to highlight the “*V* -shape” pattern.

### 3.2 A theoretical explanation for the phase transition observed in Fig. 2a

Fig. 2a exhibits a remarkable feature: there is a rather abrupt transition from effective (white) to ineffective (blue) policies, depending on the starting time of distancing policies. In this section, we examine if this observation can be understood theoretically. In other words, can we estimate the rather narrow region where the transition occurs?

We answer this question as follows. We consider the *SIR* model, where social distancing is initiated at time *t_s_*. Note that we are analyzing the behavior of the model with respect to the horizontal axis in Fig. 2; in this section we thus ignore the duration of distancing (*t_d_*) and hence assume distancing remains in effect until the end of the simulation. Here we give a characterization of peak of the infected population as a function of *t_s_*.

Fix an *SIR* model with time-varying *β*(*t*), such that

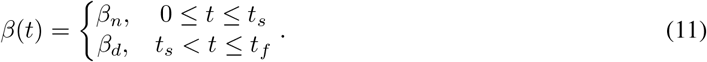

We assume that *β_n_ > β_d_*, so that distancing has the effect of reducing the transmission rate of the disease. Here *t_f_* is the entire period under consideration. Our goal is to understand the peak of the infected population, *I_p_*, as a function of the switching time *t_s_*.

Note that by changing *β_n_* to *β_d_* we have effectively reduced the basic reproduction number *R*_0_:

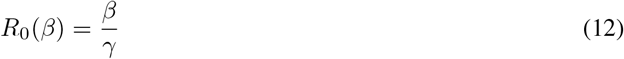

From Eq. (34), we see that such a switch then increases the value *S_p_* = *S_p_*(*β*), given by

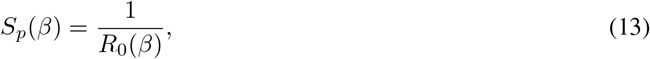

the susceptible population at the global maximum value of *I* over the *entire integral curve*

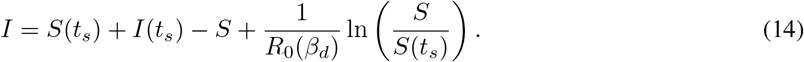

Of course, this global maximum may not be attained by the forward-time trajectory, and is generally reached if and only if the switching time *t_s_* satisfies

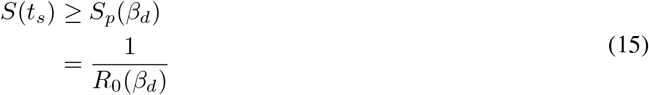

Note that if (15) is violated, the maximum value of *I* along the trajectory is given simply by *I*(*t_s_*) (see also Eq. (19) below and the subsequent discussion).

This then allows us to compute the value of *I_p_* as a function of the state (*S*(*t_s_*), *I*(*t_s_*)), where *t_s_* is the time at which social distancing is implemented. We first assume that both *β_n_* and *β_d_* are such that

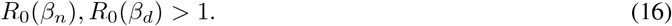

The above then implies that the infected population may increase in both the normal and distanced environments, if the susceptible population at the distancing time *t_s_* is large enough (see (19) below). Note that this matches the parameter values utilized in Fig. 2, where

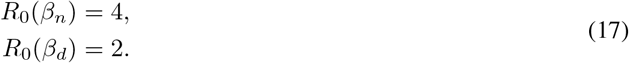

Since *R*_0_ increases as a function of *β*, Eq. (13) implies that

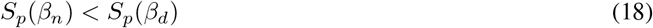

Assuming (16), we are able to compute the maximum of the infected population as a function of the susceptible population at the time *t_s_*:

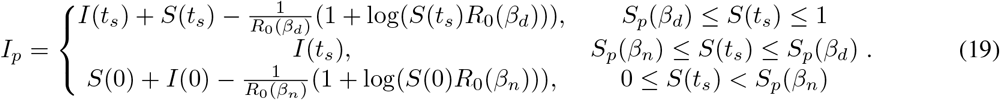

Eq. (19) can be understood as follows. Recall that *S*(0) ≈ 1 and that *S* is decreasing. Hence an early time *t_s_* corresponds to *S*(*t_s_*) close to 1, while waiting longer decreases the value *S*(*t_s_*). Hence left-to-right in Fig. 2 corresponds to top-to-bottom in Eqs. (19).

The first case occurs if *t_s_* is less than the time at which the peak value of the infective state would have been attained, if one had imposed social distancing from time zero. Note this peak always occurs, as a function of state *S*, *before* the non-distanced peak by (18). Specifically, *I* was increasing for *t* ∈ [0, *t_s_*] and at *t_s_* satisfies

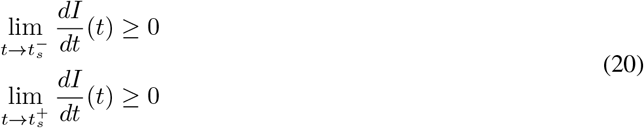

Thus, *I* is increasing at *t_s_*, and as *β*(*t*) ≡ *β_d_* for *t* > *t_s_*, the maximum of *I* (for all *t* ∈ [0, *t_f_*]) occurs at the peak of the distanced dynamics, which is given by (37), with new initial conditions (*S*(*t_s_*)*, I*(*t_s_*)) and *R*_0_ = *R*_0_(*β_d_*). This is precisely the first formula in (19).

If time *t_s_* is large enough so that *S_p_*(*β_n_*) ≤ *S*(*t_s_*) ≤ *S_p_*(*β_d_*), then *I*(*t*) must satisfy

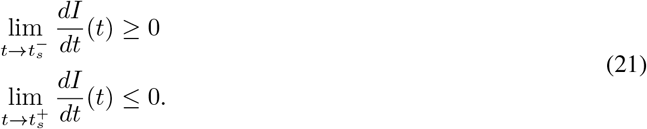

Since *I* was increasing for 0 ≤ *t* ≤ *t_s_*, the maximum of *I* occurs at *t* = *t_s_*, and hence yields the second formula in (19).

Lastly, if *t_s_* is long enough so that *S*(*t_s_*) < *S_p_*(*β_n_*), then

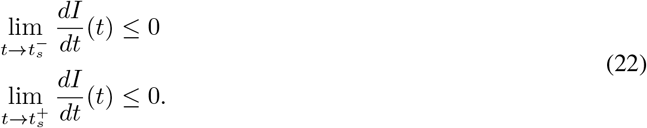

Thus, the peak of the infected compartment occurred at an earlier time, before distancing was enacted, with *β* = *β_n_*. This peak value is given precisely by (37), with *β* = *β_n_*, since the overall peak value is given by the peak value of the non-distanced dynamics; this corresponds to the third formula in (19). For a visualization of all three cases, see Fig. 4.

**Figure 4:**
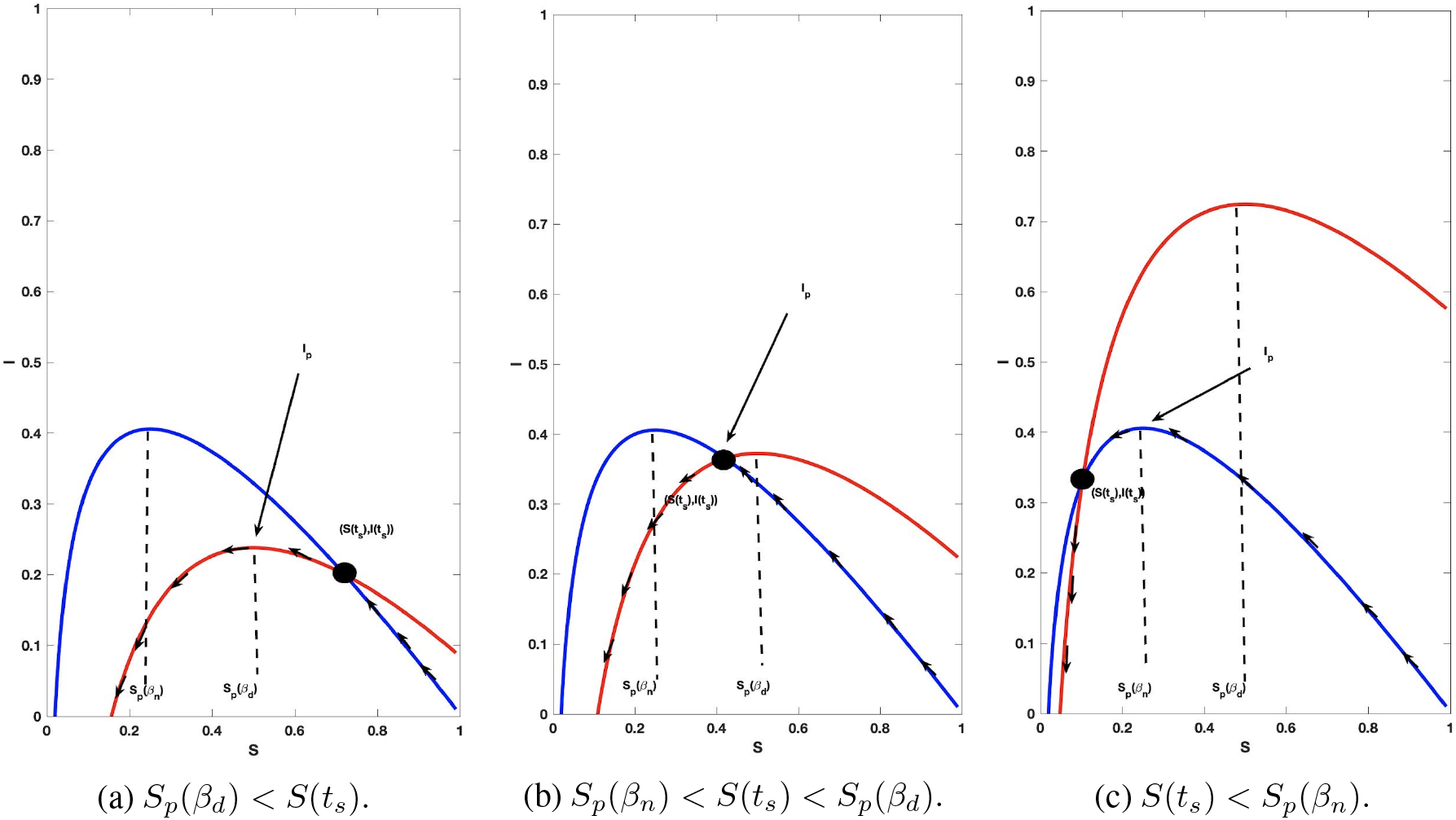
Integral curves defined by changing *β* (see Eq. (11). The blue curve corresponds to *β = β_n_* (non-distanced), and the red corresponds to *β* = *β_d_* (socially distanced). The time *t_s_* denotes the time when social distancing is enacted. All cases assume *S*(0) ≈ 1. The three figures (left-to-right) denote the three different formulas stated in Eqs. (19) for *I_p_* (top-to-bottom).

Formula (19) allows us to estimate the differential sensitivity of *I_p_* with respect to *t_s_*. First note that this sensitivity is approximately proportional to the differential sensitivity of *I_p_* with respect to *I*(*t_s_*), since (assuming *S*(0) ≈ 1 and that *S* is relatively constant on [0, *t_s_*]):

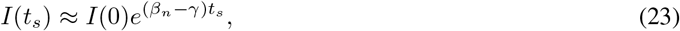

or

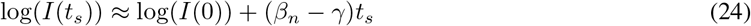

That is, the two differential sensitivities (*I_p_* with respect *t_s_*, and *I_p_* with respect to *I*(*t_s_*)) differ by a factor (*β_n_ − γ*)*t_s_*. In the remainder, we compute the differential sensitivity of *I_p_* with respect to *I*(*t_s_*) as a proxy for the true sensitivity of *I_p_* with respect to *t_s_* (the quantity of interest from Fig. 2).

We now compute the differential sensitivity of *I_p_* with respect to *I*(*t_s_*) using Eq. (19). The following analysis is for all *S*(*t_s_*) ∈ [0, 1], but we must first understand where this formula actually defines a function. *I_p_* is a function of *I*(*t_s_*) only if the latter’s range is restricted to include regions where *I*(*t_s_*) is invertible (with respect to *t_s_*). Since *I*(*t_s_*) has a relative maximum at distancing time *t_s_* such that *S*(*t_s_*) = *S_p_*(*β_n_*), it is clear that the domain of *I_p_* must be restricted into two regions: *I*(*t_s_*) with *S*(*t_s_*) ∈ [*S_p_*(*β_n_*), 1] (where *I*(*t_s_*) is non-decreasing) and *I*(*t_s_*) with *S*(*t_s_*) ∈ [0, *S_p_*(*β_n_*)) (where *I*(*t_s_*) is decreasing). With the above understanding as to an appropriate domain, we compute the differential sensitivity of *I_p_* with respect to *I*(*t_s_*):

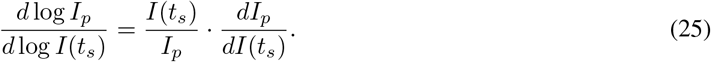

Eq. (19) then allows us to immediately compute the sensitivity if *I*(*t_s_*) is such that *S*(*t_s_*) ∈ [0, *S_p_*(*β_d_*)):

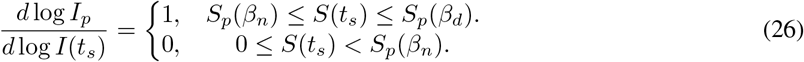

Note that the above regions correspond to Figs. 4b–4c, and is obtained by differentiating the second and third formulas in (19). In words, (26) says that the differential sensitivity is relatively large and constant for *t_s_* such that *S_p_*(*β_n_*) ≤ *S*(*t_s_*) ≤ *S_p_*(*β_d_*), and then drops to 0 for larger *t_s_* (recall that *S*(*t_s_*) decreases as a function of *t_s_*). Thus the social distancing start time should have no effect on *I_p_* if *S*(*t_s_*) < *S_p_*(*β_n_*), whereas there is a band of social distancing start times where *I_p_* increases rapidly. Note for the parameter values utilized in Figure 2, we compute

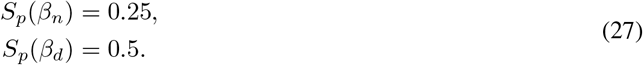

and simulating we obtain the corresponding critical start time region (where *S*(*t_s_*) = *S_p_*(*β_d_*)*, S_p_*(*β_n_*)) as

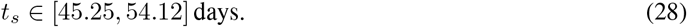

This is in close agreement with the region observed in Fig. 2, and hence describes the vertical transition band numerically computed.

Analyzing the sensitivity at earlier start times (*t_s_* such that *S*(*t_s_*) ∈ (*S_p_*(*β_d_*), 1)) is more challenging using Eq. (19), since it requires taking a derivative of *S*(*t_s_*) with respect to *I*(*t_s_*). We note that we expect the sensitivity to be small for sufficiently early distancing times *t_s_*, since we observe only slight variation in *I_p_* from Fig. 2 at small *t_s_* (and large *t_d_*, as we are not relaxing social distancing in this analysis). *Hence we conjecture that the sensitivity is largest exactly in the band given by* (28).

Numerically we compute the differential sensitivity of *I_p_* for the first two regions given in (19) (*t_s_* such that *S*(*t_s_*) ∈ [*S_p_*(*β_n_*), 1]); see the blue curve in Fig. 5. The two regions plotted correspond to the times *t_s_* where *I*(*t_s_*) is increasing. Note the approximate constant sensitivity of 1, as predicted by the first expression in (26), for *t_s_* such that *S*(*t_s_*) ∈ (*S_p_*(*β_n_*)*, S_p_*(*β_d_*)). We further observe a small sensitivity for small *I*(*t_s_*) (hence small *t_s_*), which was also expected from Fig. 2. The differential sensitivity then appears to gradually increase in value, until it reaches 1 near the transition region (*S*(*t_s_*) = *S_p_*(*β_d_*), and *I*(*t_s_*) = 0.33). As computed in (26), the sensitivity is 0 for larger *I*(*t_s_*). A small overshoot in Fig. 5 is most likely due to numerical error, as the computation requires numerical differentiation, and was pre-processed via local averaging. Hence, at least for parameter values utilized in the above, we find an interval of critical distancing start times for which the maximum of the infected population is most sensitive.

**Figure 5:**
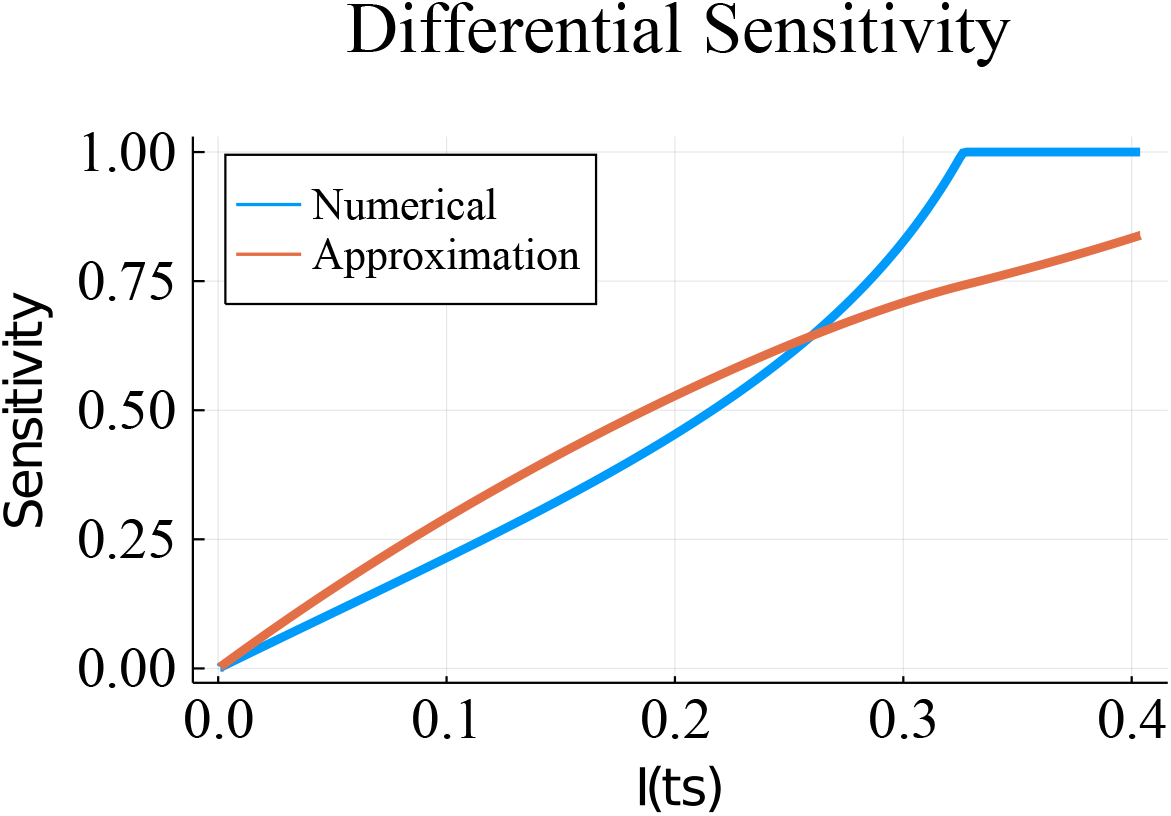
Differential sensitivity of *I_p_* with respect to *I*(*t_s_*). Numerical approximation of sensitivity is indicated by the blue curve, corresponding to region where *I*(*t_s_*) is increasing (first two formulas in (19)). *I*(*t_s_*) ∈ [0.33, 0.4] is the transition region in (19). Orange curve is approximation to sensitivity given by Eq. (30).

Lastly, we approximate the differential sensitivity in the initial region where 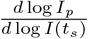 increases from 0 to 1 by making the approximation that

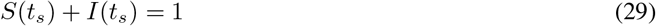

in (19). Note that this should be accurate for small *t_s_*, when *R*(*t_s_*) ≈ 0, but in general will not be true for larger times. Replacing *S*(*t_s_*) by 1 – *I*(*t_s_*) in the first formula of (19) and differentiating with respect to *I*(*t_s_*) yields the approximation

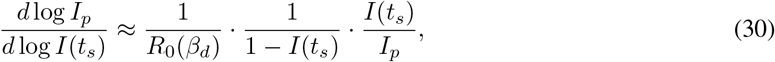

where *S_p_*(*β_d_*) < *S*(*t_s_*) < 1. This formula is the orange curve in Fig. 5. As expected, this approximation seems to be accurate initially, but soon diverges from the numerically computed sensitivity.

We make a final note concerning the above approximation related to the duration of social distancing, *t_d_*. Recall that we assumed that *t_d_* was large, and hence considered a switched system of the form (11), and thus our analysis focused on two integral curves (corresponding to *β_n_* and *β_d_*; see Fig. 4). It is possible that the reduction in transmission rate (*β_n_* → *β_d_*) is such that

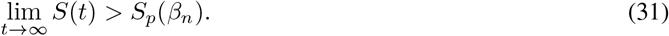

In this case, *R*_0_(*β_n_*) > 1, so that the reintroduction of even one infected individual into the population will stimulate an epidemic *if distancing mandates are relaxed* (*β_d_* → *β_n_*). Hence although the calculations are completely valid in the case of an infinitely long duration *t_d_*, the resulting dynamics in a “real-world” scenario, that is, one in which social distancing is ceased, may still result in an unstable system at risk of future outbreaks. This further necessitates the need for a careful design of single-pulse strategies of social distancing. See [57] for a further discussion of this phenomenon.

### 3.3 Are the above dynamic properties universal features of more complex models?

One may reasonably ask whether the features observed for the *SIR* model hold for more complex examples. For example, is the “*V* shape” in Fig. 2 a universal property? Perhaps surprisingly, it appears that the answer is “yes.” The *SIR* model is relatively simple, and thus its properties are easier to analyze compared to complex multi-compartment systems. However, we can use simulations for these models to investigate the universality of the phenomenon (Fig. 6).

**Figure 6:**
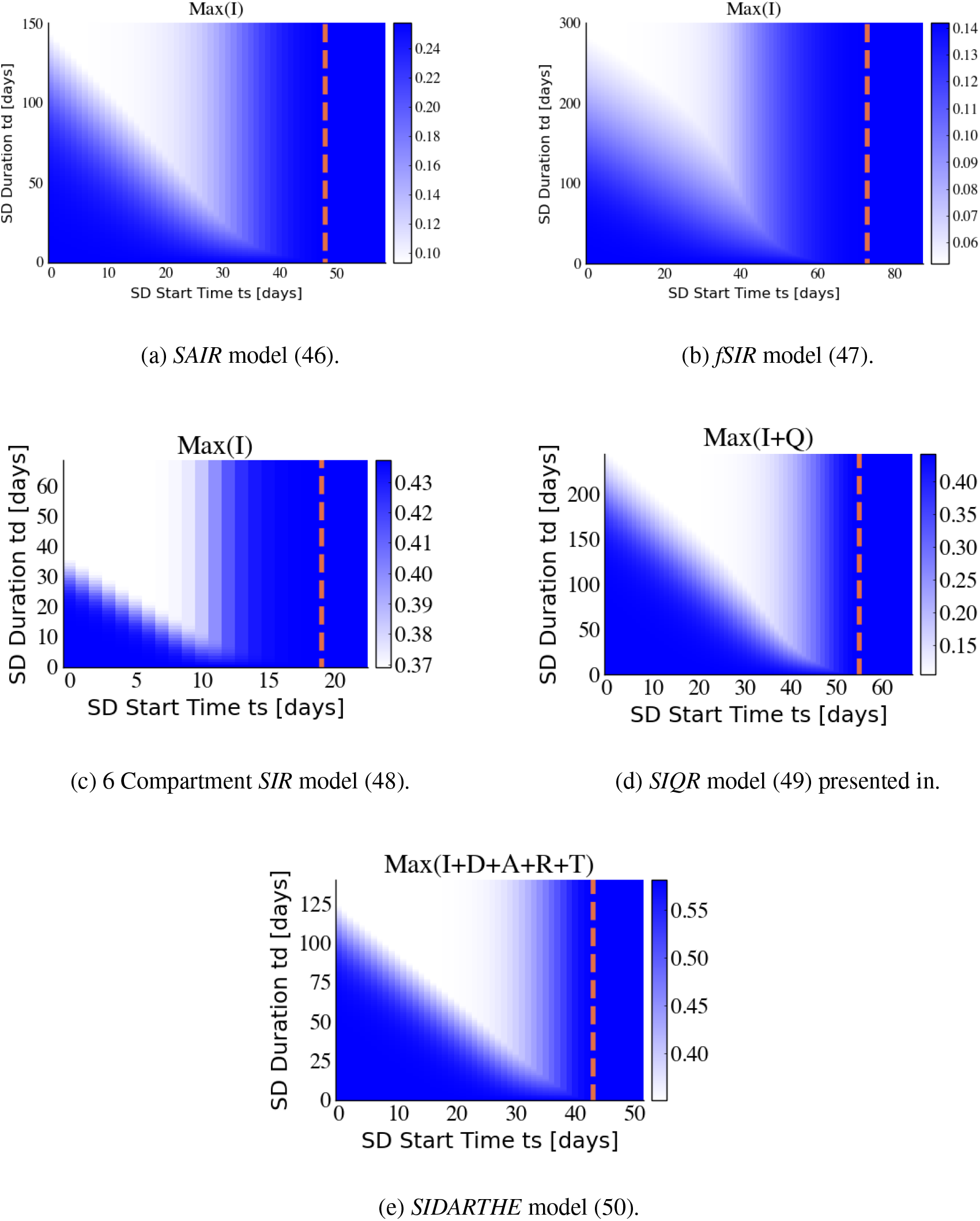
Recent models proposed to capture the COVID-19 behavior show the “V” pattern. Models and their parameters are fully presented in Appendix C. The ratio between transmission rates of normal and socially distanced populations (*β_n_/β_d_*) is fixed at 2 for all models.

We investigated several epidemic models that have been recently formulated to capture the spread of COVID-19. Each model is simulated as in Section 3.1, utilizing parameters used in the respective papers and thought to be appropriate, based on data available at the time, for the COVID-19 pandemic.

We briefly describe the models simulated in Fig. 6. The *SAIR* system is a simple variation of the *SIR* model that includes an additional compartment *A* for asymptomatic infected individuals [58, 59]. *fSIR* is an infection aware population model with the same number of compartments as the *SIR* model with the additional assumption that the contact rate between individuals decreases as a function of the infected population [39]. The 6-compartment *SIR* model is a variation of the *SIR* model that is obtained by dividing the susceptible population into two categories: socially distanced and non-distanced populations [28]. The *SIQR* model is also a variation of the *SIR* model that includes an additional compartment *Q* for the quarantined population [60], and the *SIDARTHE* model is a more complicated variation of the *SIR* model with 8 different compartments proposed to describe COVID-19 in [57].

The “V” shape pattern observed in Fig. 2 seems to be consistent among these more complicated epidemic models (Fig. 6) and similar trade-offs between the start time and duration of social distancing exist. We conjecture that this behavior is preserved for a large class *SIR*-derived models due to the both the similar network structure and control (effect of social distancing on *β*) present in many types of epidemic models.

Although the range of the infected peak is different among these models, the reduction in magnitude between the virtual peak and that under socially distancing is about 50% in every case. Also, the slope of the diagonal border between the blue (high) and white (low) regions can be approximated by Eq. (6).

### 3.4 Periodic relaxation

From an economic perspective, periodic relaxation of social distancing is favorable to prolonged single interval strategies [56]. A policy with regular periods of distancing and relaxation can significantly delay the time of the peak of the epidemic, while still allowing limited economic activity [28, 53, 56].

Fig. 7 provides numerical simulations of the *SIR* under periodic relaxation with different periods *T* (shown on the horizontal axis) and closed business-time ratios *r* (shown on the vertical axis). Here *r* is defined as the ratio between the duration of social distancing and the period of distancing. As in the previous section, the maximum infected population is plotted as a measure of the outcome of different distancing strategies. For systems that are affine in control (here viewed as the contact rate in epidemic models), fast switching policies behave similarly to *constant* social distancing regimens with transmission rate 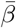:

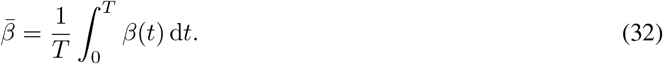

**Figure 7:**
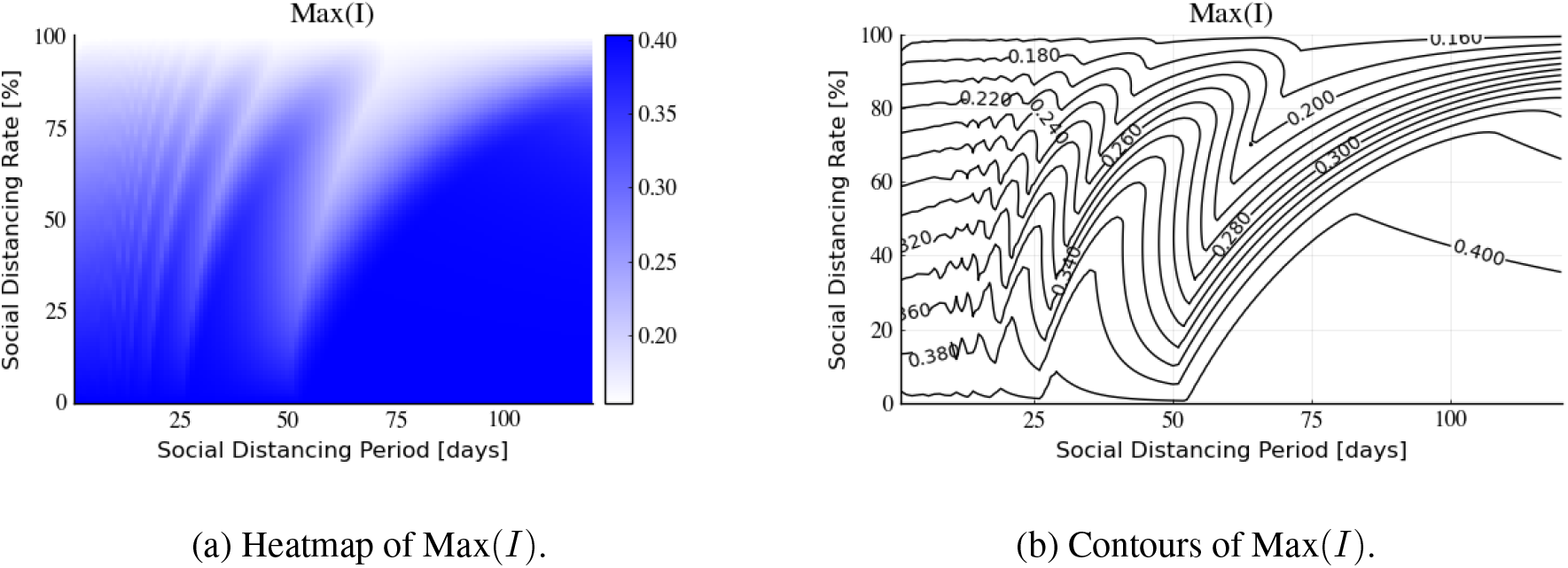
Infected peak Max(*I*) of *SIR* model (1) under various periodic distancing policies with different period *T* and ratio of social distancing *r*. See Fig. 2 for parameter values.

Recall that time *T* denotes one period of social distancing and relaxation, and hence 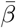 represents the time-averaged transmission rate (see Appendix B). Indeed, examining Fig. 7b, we observe that as the period *T* decreases, the infected peak converges to the response corresponding to a constant 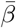 which corresponds to the value given in (32). On the other hand, as the period *T* increases, the infected peak will be less dependent on the weighted average of the transmission rate. Observe also that the infected peak is not monotonic with respect to either the period *T* of the social distancing relaxation policy or the ratio *r* of distancing. In Section 3.5, we argue that this behavior is due to “mistiming” the social distancing mandate. This behavior has been also observed in [28, 53].

The limitation of such policies is that the peak of the infected population is a function of both *β_n_* and *β_d_*, while the peak of the infected population with a single interval of social distancing at a *proper time* will only be dependent on the social distancing transmission rate *β_d_*. This can be seen in Fig. 9, where we observe that the peak of the infected population during a single interval social distancing strategy is almost identical to the infected peak for a population that is indefinitely distanced (compare the maximums of the solid blue and black curves). In what follows, we propose a combination of a single-pulse and periodic social distancing policy as an approach that optimally reduces the infected peak while both allowing periodic economic activity and delaying the onset of the infected peak.

### 3.5 Periodic relaxation combined with a single interval of social distancing

The peak of infected population depends non-monotonically on both the period *T* and ratio *r* of periodic policies. A periodic social distancing relaxation policy with a large period time *T* may lead to high transmission rates at the critical time of an epidemic (e.g. at the potential peak of the infected population). Or it may lead to a well-timed strategy and hence significantly reduce the infected peak. To address this uncertainty, we propose combining single-pulse social distancing with periodic relaxation (Fig. 8).

**Figure 8:**
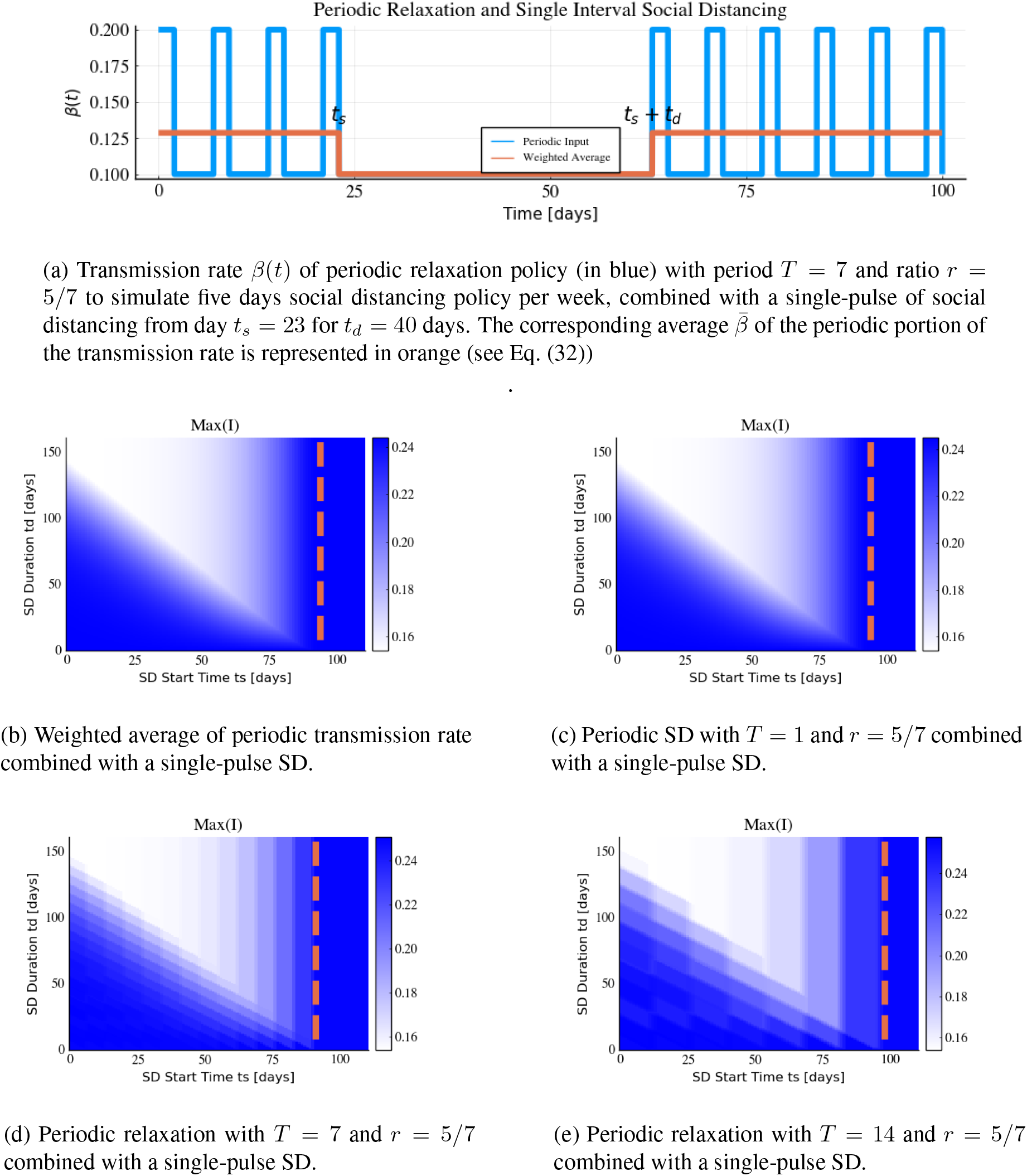
Effect of periodic relaxation policy in combination with a single interval social distancing for *SIR* model with parameters *β_n_* = 0.2, *β_d_* = 0.1, and *γ* = 0.05. The dashed orange line, which represents the infected compartment peak time without single interval social distancing policy (virtual peak), is represented at *t_s_* = 93.78, *t_s_* = 94.0, *t_s_* = 91.0, and *t_s_* = 98.0 in Figs. 8b, 8c, 8d, and 8e, respectively.

Fig. 8 illustrates the effect of combining a single pulse of social distancing and periodic relaxation policies for different periods *T* in the *SIR* model. The transmission rate used as an input for the pulsed strategy is visualized in Fig. 8a. This representation is for periodic social distancing with period of one week (*T* = 7 days), with two days of relaxation and five days of distancing enacted, i.e. *r* = 5/7. The single-pulsed strategy with different start time *t_s_* and duration *t_d_* is simulated in combination with a weighted average transmission rate (Fig. 8b) and periodic pulsed strategies with different periods of *T* = 1 day, *T* = 7 days, and *T* = 14 days in Figs. 8c, 8d, and 8e, respectively. Note that for small period *T* = 1, the periodic relaxation policy is well-approximated by the average transmission rate as discussed in Section 3.4; compare Figs. 8b and 8c. However, for larger *T*, the corresponding simulations differ significantly (e.g. Figs. 8b and 8e).

We first note that the single interval of *strict* social distancing may generally be delayed for mandates which include periodic relaxation; compare the horizontal axes (*t_s_*) in Figs. 2 and 8. This is advantageous, as it allows for a longer amount of time for medical professionals and policy-makers to prepare for the peak of the epidemic. Furthermore, the amplitude of the infected peak is lower when a periodic relaxation policy is combined with a single interval social distancing. By comparing the dark blue (high) and white (low) regions in Fig. 8 it can be observed that the infected peak can be reduced by 30% percent with just one additional single-pulse social distancing at an appropriate time. The “V” shape pattern is still consistent for the combination of periodic relaxation and signal interval social distancing policies. Such policies are feasible to implement during an epidemic, and still provide opportunities for limited economic activity. Hence a *well-timed* additional single-pulse of strict social distancing utilized in conjunction with periodic relaxation may provide an appealing response to an epidemic.

## 4 Conclusions and Discussion

The white area of the “V” shaped graphs shown in Figs. 2a and 6 represents a linear trade-off between the start time *t_s_* and the duration *t_d_* of social distancing. This phenomenon has been illustrated for several different epidemic models, some of which were recently proposed for COVID-19. A single interval social distancing would not be effective in the reduction of the infected peak if the start time is too late (blue region on the right side of vertical line), or too early (the blue area on the bottom left side of the diagonal line). On the other hand, a single interval of social distancing enacted at a *proper time* can significantly reduce the infected peak. Combining such a single interval with periodic relaxation also appears to significantly reduce the peak of the infected population, while simultaneously delaying the necessary start time of the strict distancing period, and hence provides an appealing option for policy-makers.

### 4.1 Comparison of optimal single shot and complete social distancing strategies

Fig. 9 is an example of how *well-timed* social distancing can reduce disease burden for each of the investigated models. The dashed black line is the time-varying transmission rate *β*(*t*), defined by (3), modeling a single interval of social distancing. The solid black curves represents the response of the infected compartment to this *β*(*t*), while the red and blue curves show the response to no and full social distancing, respectively. That is, comparing to (3), the red curves correspond to a system with *β*(*t*) ≡ *β_n_*, while the blue curves correspond to *β*(*t*) ≡ *β_d_*. We see that the values of the peaks of the black and blue lines are similar, implying that *properly* timing a single pulse of social distancing can have a significant impact on the infected peak; indeed, the response to such regimen appears to approximate the response to a fully isolated distancing policy (blue curves).

**Figure 9:**
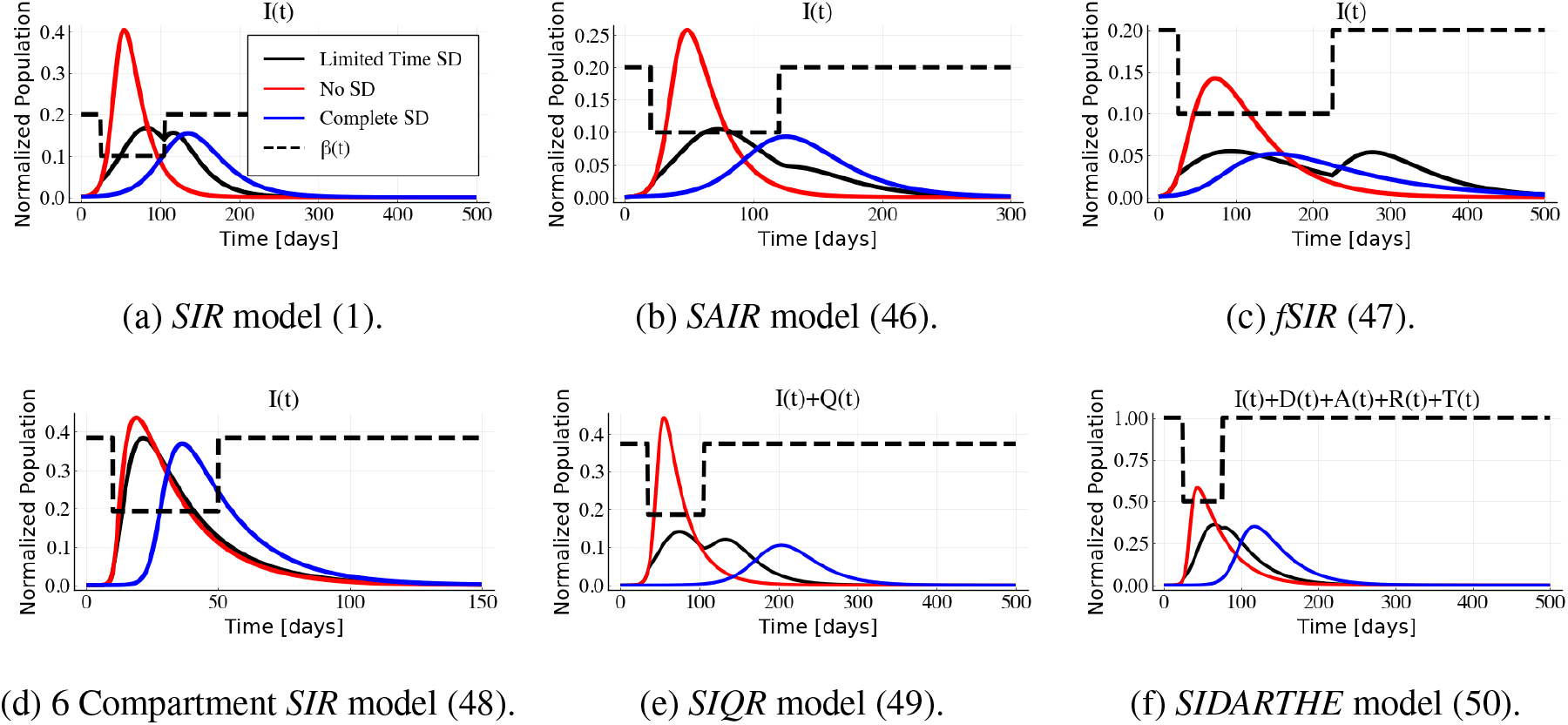
Optimal social distancing and quarantine period during an epidemic based on different models: a) The *SIR* model (1), b) the *SAIR* model presented in Section C.2, c) the six-compartment *SIR* model with social distancing from [28] presented in Section C.4, d) the *fSIR* [39] presented in Section C.3, e) the *SIQR* model [60] presented in Section C.5, and f) and the *SIDARTHE* model [57] presented in Section C.6. Dashed line shows the input *β*(*t*) applied in each model as a social distancing control. The black line represents the infected compartment with limited time social distancing, the red line shows the dynamics of the infected compartment with *β*(*t*) = *β_n_* (no social distancing), and the blue line represents the scenario of having the social distancing during an entire epidemic with *β*(*t*) = *β_d_*.

Based on Figs. 6 and 2, the start time *t_s_* and the duration *t_d_* of a single interval of social distancing which minimize the infected peak are not unique. Different values of *t_s_* and *t_d_* that correspond to the white color in the heat maps/contour plots will reduce the infected peak to approximately the same degree. Furthermore, if we define the optimal interval of social distancing start time based on a given social distancing duration, then the range of acceptable social distancing start times increases as the social distancing duration increases. For example, examining Fig. 2a, we see that a social distancing policy with a duration of *t_d_* = 75 days should be commenced at approximately *t_s_* ∈ [25, 30], while a distancing policy with a longer duration of *t_d_* = 100 days should be commenced at approximately *t_s_* ∈ [20, 30]. This latter observation is important from an implementation perspective, as a larger range of start times implies less precision is necessary for designing social distancing strategies. As parameter estimation is difficult, especially in an evolving pandemic (e.g. March 2020), more robust protocols may be desirable.

### 4.2 Robustness of “V” shape pattern

The ratio between transmission rates, *β_n_/β_d_*, is set to 2 for the simulations represented in Figs. 2, 6, 7, 8, and 9. This ratio is seen to be approximately proportional in magnitude to the slope of the diagonal border between the blue and white regions of Fig. 2 in Section 3.1 (see (6) for a more precise expression). By increasing the transmission rate ratio *β_n_/β_d_*, i.e. by introducing a more draconian social distancing policy, the magnitude of the slope of the diagonal line increases. Fig. 10 shows the heat map of the normalized infected peak of the investigated models when, instead, *β_n_/β_d_* = 10. Note that *β_n_* is the same for the investigated models in Figs. 6 and 10. It can be observed that social distancing pulses that take place at an early time of an epidemic will not be effective in diminishing the peak of the infected population. Intuitively, the blue area on the left side of vertical white region is for the scenarios when the infected peak occurs after the social distancing time interval. But, if a social distancing is imposed from the white region based on the suggested start time *t_s_* and duration *t_d_* of social distancing, then it will be very effective in inhibiting the peak of the infected population. The persistence of the characteristic “V” shape and its close agreement with the predictions presented in Sections 3.1 and 3.2 show that our analysis is robust with respect to parameter variations.

**Figure 10:**
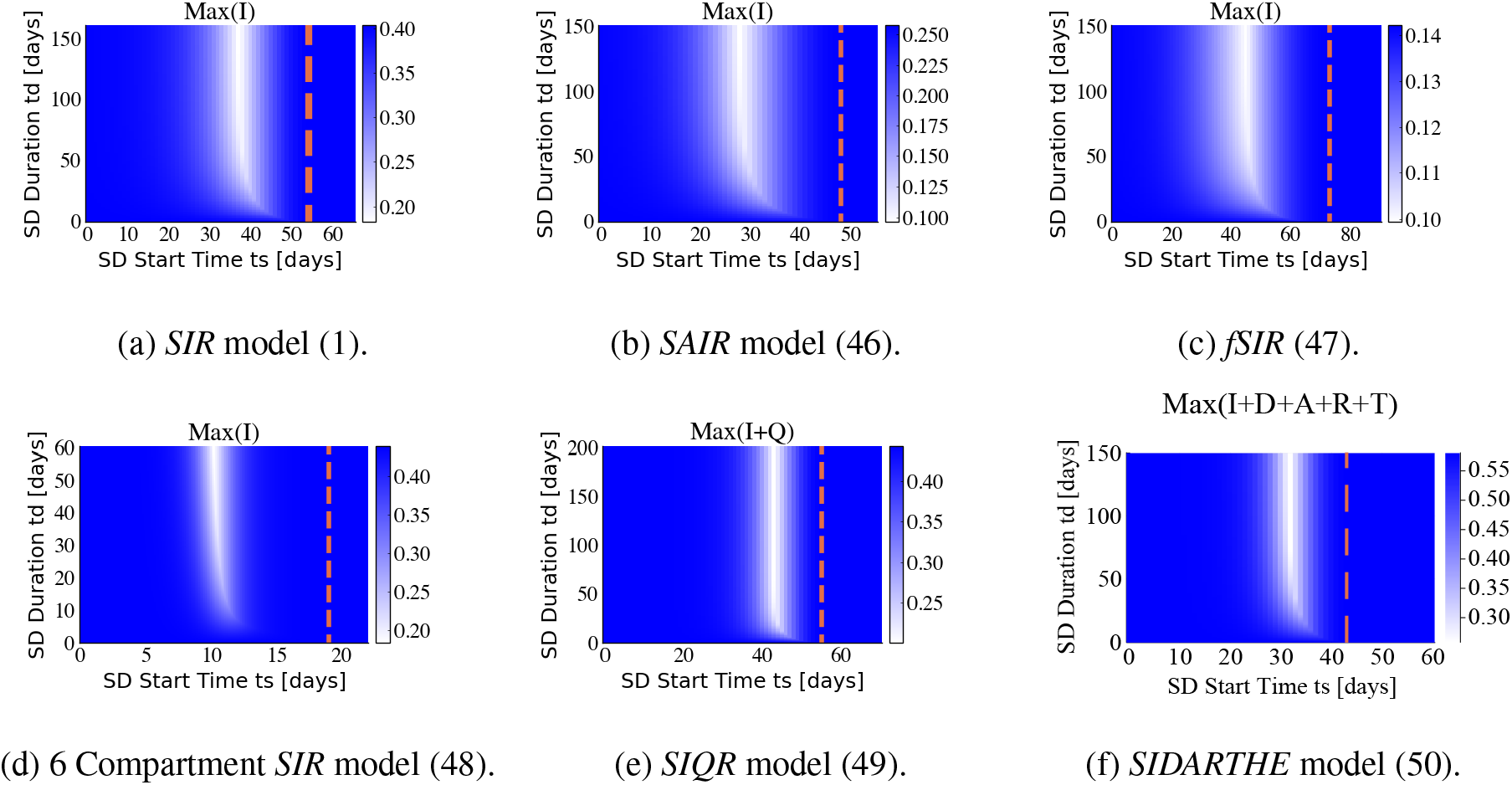
Maximum of the infected compartment for a selection of models introduced to capture the dynamics of the COVID-19 pandemic. Models and their parameters are fully presented in Appendix C. The ratio between transmission rates in the single-pulse strategy is *β_n_/β_d_* = 10 (see equation (3))

### 4.3 Discussion

In this work we have investigated the response of epidemic models to social distancing as a control strategy. We have analyzed the basic features of the classical *SIR* model with respect to a single interval of social distancing both computationally and theoretically. We have shown, via computations, how the basic response structure of the *SIR* model is preserved for a number of more complex (but related) epidemic models. Importantly, although a strict social distancing mandate enacted early on in an epidemic will delay and potentially reduce the peak of the infection, too early relaxation may inhibit a significant reduction. Similarly, waiting too long to implement distancing will also be ineffective as a mitigation strategy, but importantly, this critical delay occurs earlier than expected, and can be quantified.

More precisely, we have introduced a trade-off between the start time and the duration of social distancing effects on the peak of the infected population. A “V” shape pattern has been observed for the *SIR* model. This pattern can be understood mathematically in the special case of the *SIR* model and is verified through simulation for more complex models recently proposed for COVID-19. A single pulse of social distancing is shown to be most effective if it happens at a *proper time*. Moreover, the infected peak for economically preferable strategies like periodic social distancing relaxation can be reduced by a single pulse of social distancing.

## Computational Resources

The numerical simulations and plots are done with the DifferentialEquations package [61] of Julia programming language [62], and the analysis notebook are available on https://github.com/sontaglab/epidemics repository.

## Acknowledgments

We thank M. Ali Al-Radhawi for his comments. We thank the anonymous referees for a large number of very constructive suggestions, which much improved the manuscript.

## A *SIR* Model

### A.1 Peak of the infected compartment

The peak of the infected compartment for the simple *SIR* model can be characterized at the time when 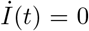, from Eq. (1). This is the time when the rates at which populations are being infected and recovered (or removed) from the infected compartment balance out. We will denote the peak of *I* by *I_p_*, and the corresponding susceptible population by *S_p_* and we write

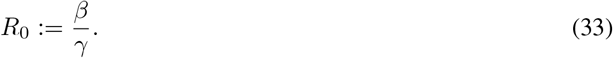

for the basic reproduction number of the *SIR* model. Provided that 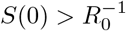, one has:

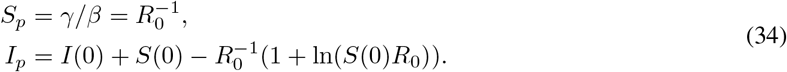

Note that the maximum of the infected compartment is only a function of the basic reproduction number *R*_0_. The formula for *S_p_* is immediate from 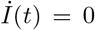. The formula for *I_p_* can be derived as follows. We have the following implicit formula between the susceptible *S* and infected *I* compartments:

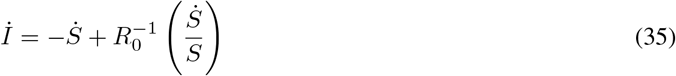

which implies

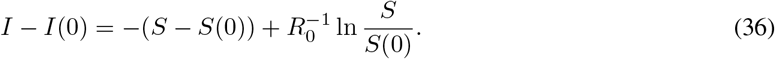

Here *S*(0) and *I*(0) are the initial values of the susceptible and infected compartments at the beginning of the epidemic. Therefore, the peak of the infected compartment is:

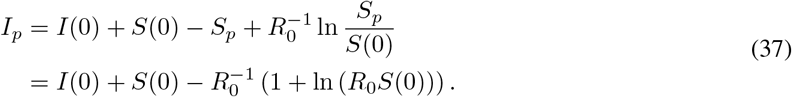

At the start of the epidemic, *S*(0) ≈ 1 and *I*(0) ≈ 0, so the expression for the infected peak is simplified to:

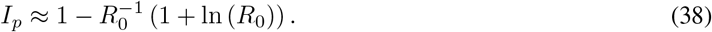

### A.2 Change of variables for *SIR* model

To gain a better understanding of the infected compartment peak time *t_p_*, we can reparametrize the time *t* by *τ* in the form of d*τ*/d*t* = *I*(*t*) with initial time *t*_0_ = *τ*_0_ = 0 as suggested by [63], the *SIR* model(1), under this change of time scale, is now linear:

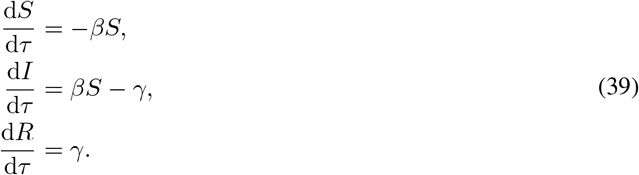

We remark that his transformed system evolves from *τ* = 0 to 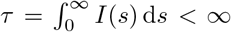. The solution of the linearized model can be written in the following form

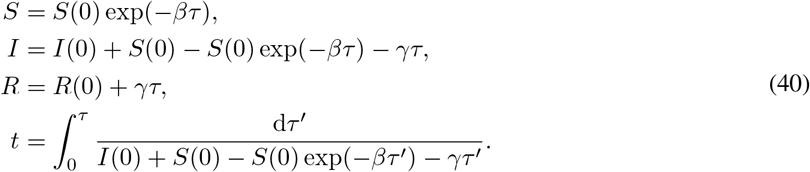

The infected peak and its corresponding time can be written analytically as follows:

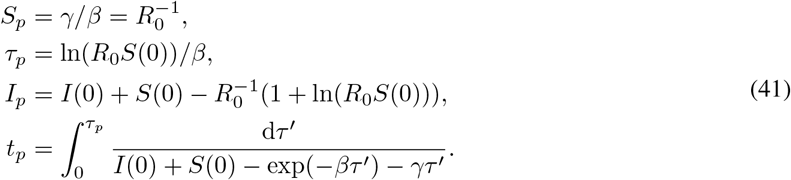

It can be observed that the maximum of the infected compartment is only a function of the basic reproduction number *R*_0_, while the time of the peak *τ_p_* is dependent on *R*_0_ and *β*. We remark that a different time scaling and shifting have been used by [64] to show statistical similarities of the COVID-19 spread in different countries.

### A.3 Linear approximation

The integral in Eq. (40) is hard to compute explicitly. To obtain an intuitive approximation for small times for *I*(*t*) from Eq. (39), we’ll consider *S*(*t*) to be constant, and therefore equal to its initial value *S*(0). That gives:

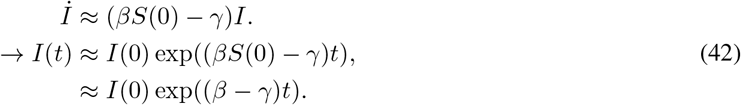

This approximation is reasonable for the beginning of the spread of the virus, when *S*(*t*) is close to one, the total population. We used this approximation in Section 3.1.

## B Fast Switching Policies

Consider a general system affine in controls ([65]):

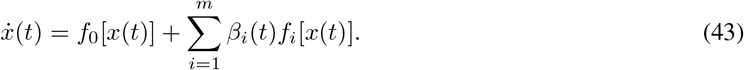

We justify here the fast-switching remark made earlier. We use the general system affine in controls formalism in Eq. (43). Let us suppose that the inputs *β_i_*(*t*) are periodic with period *T*, and consider the constant control obtained by averaging the *β_i_* over a period:

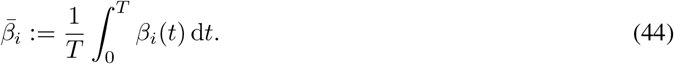

It can be proved that, if the frequency of *β*(*t*) = (*β*_1_(*t*),…, *β_m_*(*t*)) goes to infinity, meaning that the switching time approaches 0, then solutions approach those for the average 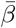. This follows from standard averaging results for systems that are affine in controls. Specifically, (1) the map from controls on an interval [0, *T*] to trajectories on [0, *T*] is continuous with respect to the weak* topology in *L*^1^ and the uniform topology on continuous functions, respectively (see, e.g. [65], Theorem 1), and (2) for a periodic input *u*(*t*), the input *u*(*ωt*) converges weakly to the average of *u* as *ω* → ∞. An alternative proof can be found in [66] (Section 10.2) (changing time scale in the statement of Theorem 10.4, by *x*(*t*) = *x*(*t/ϵ*)). (This fact was also observed in [53].)

## C State space representation of several epidemic models

In order to describe several epidemic models that have recently proposed for COVID-19, we use the standard control theory state space formalism (e.g. [65]) for systems that are affine in controls *β_i_*(*t*) of Eq. (43). When there is just one control (*m* = 1), we write simply *β* instead of *β*_1_.

### C.1 *SIR* model

The *SIR* model represented in (1) can expressed, using the state *x* = (*S, I, R*), in the following form:

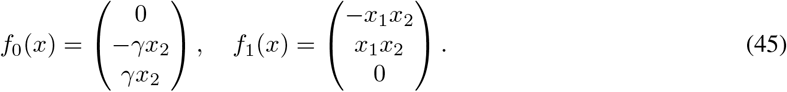

The transmission rate *β* and recovery rate *γ* are dependent on the nature of the epidemic disease, and medical resources available at the community [67–69]. From the acceptable range of parameters we used (*β, γ*) = (0.2, 0.05) and initial conditions of (*S, I, R*)_*t*=0_ = (1 − 10^−3^, 10^−3^, 0) in all numerical simulations. The value of *β_d_* used for the social distancing intervals of Figs. 2, 7, 8, and 9 is 0.1, and 0.02 for Fig. 10a.

### C.2 *SAIR* model

The *SAIR* model as a simple extension of the *SIR* model with an asymptomatic *A* compartment in the population [58, 59]. The state space representation in terms of the state *x* = (*S, A, I, R*) is:

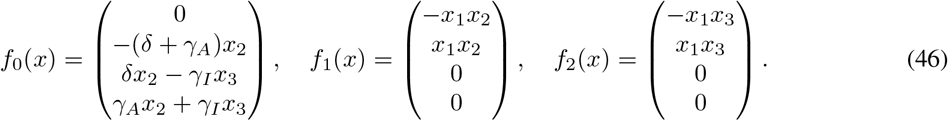

We use the parameters (*β*_1_, *β*_2_, *δ, γ_A_, γ_I_*) = (0.3, 0.2, 0.1, 0.09, 0.05) (adopted from [59]) with initial conditions of (*S, A, I, R*)_*t*=0_ = (1 − 1.1 × 10^−3^, 10^−3^, 10^−4^, 0). The values of *β*_1_(*t*) and *β*_2_(*t*) used for the social distancing intervals of Fig. 6a and Fig. 10b are half (*β_n_/β_d_* = 2), and 10% (*β_n_/β_d_* = 10) of their original value respectively.

### C.3 *fSIR* model

An“infection aware” distancing model has been recently introduced by [39], in order to account for how individuals practice enhanced social distancing as the number of infections increases. This model has an additional term of the form 1*/*(1 + *kI*) in comparison with the simple *SIR* model (1) where *k* is the feedback gain of the statistical social awareness effect on disease spread. The state variable is *x* = (*S, I, R*) and the state space representation is as follows:

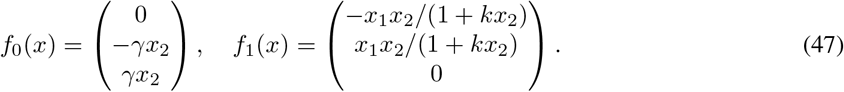

We use the parameters (*β, γ, k*) = (0.2, 0.05, 10) (adopted from the acceptable range of parameters defined by [39]) with initial conditions are (*S, I, R*)_*t*=0_ = (1 − 10^−3^, 10^−3^, 0). The values of *β*(*t*) used for the social distancing intervals of Fig. 6b and Fig. 10c are 0.1, and 0.02 respectively.

### C.4 Six-compartment *SIR* model

The 6 compartment *SIR* model [28] with state variable *x* = (*S_D_, S_N_, A_D_, A_N_, I_D_, I_N_, R*) is:

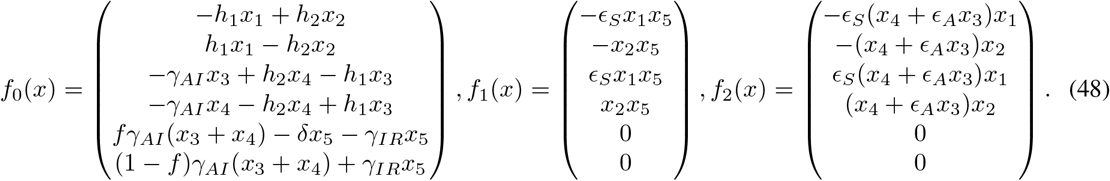

The parameters (*β_A_, ∈_A_, ∈_S_, β_I_, h_2_, γ_AI_, γ_AR_, f, δ*) = (1.4, 0.12, 0.12, 0.336, 0, 0.296, 0.048, 0.65, 0.0024) and initial conditions (*S_D_, S_N_, A_D_, A_N_, I_D_, I_N_, R*)_*t*=0_ = (0, 1 − 10^−5^, 0, 0, 10^−5^, 0), are used in our numerical simulations (as suggested by the authors [28]). The values of *β*_1_(*t*) = *β_A_*(*t*) and *β*_2_(*t*) = *β_I_* (2) used for the social distancing intervals of Fig. 6c and Fig. 10d are half (*β_n_/β_d_* = 2), and 10% (*β_n_/β_d_* = 10) of their original value respectively.

### C.5 *SIQR* model

The *SIQR* model [53, 60] with state variable *x* = (*S, I, Q, R*) is:

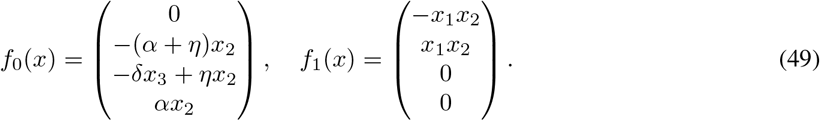

The parameters (*β, α, η, δ, N*) = (0.373, 0.067, 0.067, 0.036, 10^7^) and initial conditions (*S, I, R*)_*t*=0_ = (10^7^ − 83.333, 83.333, 0, 0) are used based on the authors suggestion [53]. The value of *β*(*t*) = *qβ* used for the social distancing intervals of Fig. 6d and Fig. 10e is half (*β_n_/β_d_* = 2), and 10% (*β_n_/β_d_* = 10) of their original value respectively.

### C.6 *SIDARTHE* model

The *SIDARTHE* model [53, 57] with state variable *x* = (*S, I, D, A, R, T, H, E*) is:

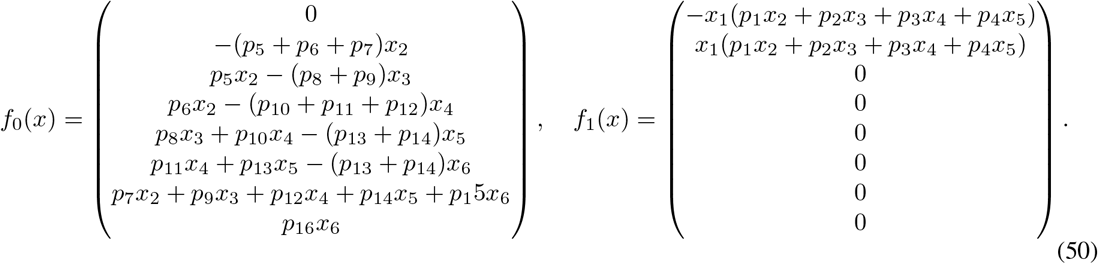

The parameters are *N* = 10^7^ and the *p_i_* represented in the following and initial conditions (*S, I, D, A, R, T, H, E*)_*t*=0_ = (10^7^ − 83.333, 83.333, 0, 0, 0, 0, 0, 0) are used for the numerical simulations based on [53]. The value of *β*(*t*) = *qβ* used for the social distancing intervals of Fig. 6d and Fig. 10e are half (*β_n_/β_d_* = 2), and 10% (*β_n_/β_d_* = 10) of their original value respectively. The parameters *p_i_* are the entries of the following vector *p*:

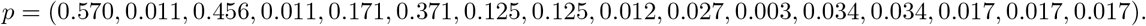

## D Infections approach zero asymptotically

We remark that, in all the models discussed, the infective compartments approach zero as time *t* → ∞. For epidemics models with no controls, this simple fact is typically established by appealing to the LaSalle Invariance Principle. However, the LaSalle principle does not apply to time-varying systems, which are the focus of this paper, so a slightly more complicated argument is needed. We illustrate with the six-compartment *SIR* model from [28], discussed in Section C.4, which we repeat here for convenience:

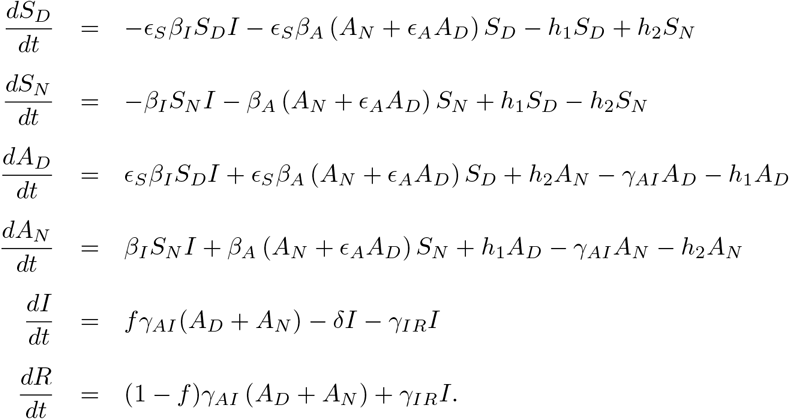

We assume that the inputs (*β* functions) are Lebesgue measurable and essentially bounded, which is a very mild assumption and covers all piecewise continuous bounded inputs. An important remark, which will be applied in the proof, is as follows. For any system *dx/dt* = *f* (*x*(*t*), *u*(*t*)) for which inputs are bounded, and for trajectories that stay in a compact set (our variables are nonnegative and add to ≤ 1), the solution *x*(*t*) is uniformly continuous. (Assuming that *f* is continuous.) This is proved as follows. In general, trajectories of control systems are locally absolutely continuous (see for example Appendix C in [65]). Thus, they have derivatives almost everywhere, and satisfy 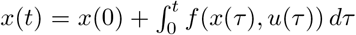 for all *t*. Therefore, the norm of the solution satisfies |*x*(*t*) − *x*(*s*)| ≤ *c|t − s|*, where *c* is an upper bound on |*f* (*x*(*t*), *u*(*t*))|. This proves that *x* is globally Lipschitz, and thus in particular uniformly continuous. In particular, of course, each coordinate of *x* is also uniformly continuous.

We will show that all the infective populations, *A_N_* (*t*), *A_D_*(*t*), and *I*(*t*) converge to zero as *t* → ∞. The paper [28] does the proof in the case of constant inputs. As there, we start by considering the total population fraction *N* defined by *N*:= *S_D_* + *S_N_* + *A_D_* + *A_N_* + *I* + *R* and observe that *dN/dt* = −*δI* ≤ 0.

The function *N* is non-increasing (because *I* is nonnegative) and thus converges monotonically to some limit. Since *dN/dt* is a constant multiple of *I*, which is uniformaly continuous (see previous remark), also *dN/dt* is uniformly continuous. Barbarat’s Lemma [66] says that a differentiable function (in this case, *N*) with a limit as *t* → ∞, and whose derivative is uniformly continuous, has a derivative that converges to zero. Therefore *I* = (1*/δ*)*dN/dt* also converges to zero.

Next we want to prove that the remaining infective compartments also tend to zero. If we prove that *dI/dt* converges to zero, then since *I* converges to zero and *fγ_AI_* (*A_D_* + *A_N_*) = *dI/dt* + (*δ* + *γ_IR_*)*I* and since both *A_D_* and *A_N_* are nonnegative, we can conclude that both *A_D_*(*t*) and *A_N_* (*t*) converge to zero, as claimed. It remains to prove that *dI/dt* converges to zero. For this, we will again apply Barbalat’s Lemma, this time to *I*(*t*). We already know that *I*(*t*) converges, so all that remains to check is that its derivative is uniformly continuous, which follows, again because *dI/dt* = *fγ_AI_* (*A_D_* + *A_N_*) − *δI* − *γ_IR_I*, from the fact that both *A_D_* and *A_N_* are uniformly continuous. This completes the proof.

## Notes

### Competing Interest Statement

The authors have declared no competing interest.

### Summary of Updates

Several edits made in response to referee comments; fixed some small errors; redid some figures.

